# Using a novel web application for thermodynamic characterization of *Tel22* G-quadruplex unfolding

**DOI:** 10.1101/730432

**Authors:** I. Prislan, S. Sajko, N. Poklar Ulrih, L. Fuerst

**Affiliations:** University of Ljubljana, Biotechnical Faculty; University of Ljubljana, Faculty for Chemistry and Chemical Technology; University of Ljubljana; University of Ljubljana, Faculty of Computer and Information Science

## Abstract

Measuring and quantifying thermodynamic parameters that determine stability of and interactions between biological macromolecules is an essential and necessary complement to structural studies. Although several laboratories are able to obtain basic thermodynamic parameters for the observed process, the data interpretation and analysis quality of reported data can be extremely tedious. We have started to develop a web application that will help users to perform thermodynamic characterization of G-quadruplex unfolding. The application can perform global fitting of calorimetric and spectroscopic data, and it uses a three-state equilibrium model to obtain thermodynamic parameters for each transition step: the Gibbs energy, the enthalpy, and the heat capacity. As well as these, the application can define the number of K^+^ ions and the number of water molecules being released or taken up during the unfolding. To test our application, we used UV spectroscopy, circular dichroism, and differential scanning calorimetry, to monitor folding and unfolding of a model 22-nucleotide-long sequence of human 3’-telomeric overhang, known as *Tel22*. The obtained data was fed to the web application and global fit revealed that unfolding of *Tel22* involves at least one intermediate state, and that K^+^ ions are released during the unfolding, whereas water molecules are taken up.

**STATEMENT OF SIGNIFICANCE:** The laws of thermodynamics provide tools for the use of elegant mathematical expressions to describe stabilities and interactions of biological macromolecule. Even though thermodynamic profiles of simple transitions (e.g., two state) can be obtained in a relatively straightforward manner, performing thermodynamic analysis of complex/ multistep transitions or global analysis of several experimental data requires some experiences and skills. In the present study we are demonstrating how newly developed web application can be used to provide better understanding of driving forces responsible for the structural interconversion of G-quadruplex structures. We have tested this web application with experimental data obtained from monitoring thermal folding/ unfolding of the 5’-AG_3_T_2_AG_3_T_2_AG_3_T_2_AG_3_-3’ (*Tel22*) DNA sequence. We believe that this application can be used as a research and/or teaching tool, and it will allow comparisons of the thermodynamic parameters obtained between different laboratories.

## INTRODUCTION

The laws of thermodynamics provide tools for the use of elegant mathematical expressions to describe stabilities and interactions of biological macromolecules. Although only a handful of people can follow what experts in thermodynamics are talking about, or whether what they are saying is relevant, thermodynamic studies have fought their way to become the essential and necessary complement to structural studies, to fully define the driving forces of folding and binding interactions (1). Accessibility of modern instrumentation to calorimetry means that more laboratories have the potential to measure enthalpy values for binding reactions and conformational transitions of biomolecules. Complete thermodynamic profiles of simple transitions (e.g., two state) can now be obtained in a relatively straightforward manner using software solutions provided by instrument manufactures. However, if thermodynamic analysis requires implementation of complex/ multistep transitions or global analysis of experimental data obtained from several experimental techniques (2), researchers have no other option but to develop their own solutions. As few have the skills or time to build tailor-made software, commercial programs are used relatively indiscriminately for different systems, which can prevent meaningful in-depth thermodynamic analyses, or – even worse – can lead to wrong conclusions.

Based on our experience of thermodynamic analysis of biological systems (3-6), we are developing a web application that can be accessible to everybody, and that will allow easy thermodynamic analysis of calorimetric and spectroscopic data. This application can be used as a research and/or teaching tool, and it allows comparisons of the thermodynamic parameters obtained between different laboratories. We have tested this web application with experimental data obtained from monitoring thermal folding/ unfolding of the 5’-AG_3_T_2_AG_3_T_2_AG_3_T_2_AG_3_-3’ (*Tel22*) DNA sequence.

*Tel22* represents a model sequence for human telomeric repeats of d(TTAGGG), and it is rich in guanines. Sequences that are rich in guanines can form four-stranded structures known as G-quadruplexes. These are composed of several layers of G-quartets, where four guanines are associated in planar tetrades. G-quadruplexes are not held together by classic Watson-Crick base pairing, but by Hoogsteen base pairing (7, 8). Although the configuration of G-quadruplexes might be intramolecular or intermolecular (e.g., bimolecular, tetramolecular) (9), it has been proposed that intramolecular G-quadruplexes are more relevant *in vivo* (10).

Blackburn and Gall studied macronucleus DNA of the ciliate protozoan *Tethrahymena thermophila*, and they showed that chromosome ends consist of tandem repeats of guanine-rich sequences (11). This was the beginning of the path that led to the discovery of telomerase (12) and the sequence of the human telomere, which consists of tandem repeats of d(TTAGGG) (13, 14). The formation of stable G-quadruplexes in the region of telomeric single-stranded overhangs has been shown to inhibit telomerase activity (15-17), and this has thus kindled an interest in telomeric G-quadruplexes as promising targets for anticancer agents (18-21). Such agents should have exquisite specificity for G-quadruplexes, but defining them requires an understanding of the driving forces that are responsible for the structural interconversions of several G-quadruplex structures. These forces are strictly connected to the solution conditions; i.e., temperature, cation type, and water activity, which have major roles in the formation of G-quadruplexes and in their physicochemical properties (22-26).

G-quadruplex structures can fold into a variety of topologies (27). Nuclear magnetic resonance and some biophysical techniques have been used to show that at room temperature and in the presence of K^+^ ions, *Tel22* can adopt different structures (28, 29). Indeed, it has been suggested that in K^+^ solutions, *Tel22* appears as a mixture of two different intramolecular (3+1) G-quadruplexes (*Hybrid-1, Hybrid-2*) (30, 31). While the *Tel22* structures have been well characterized, their folding pathways are not yet fully understood. On the basis of *in-silico* studies (32-34), it has been suggested that triplex intermediates can participate in the folding of the human telomeric quadruplex, which has been confirmed to a certain extend by *in-vitro* studies (3, 35). However, the unfolding of quadruplex structures still has not revealed all its secrets. Studying and understanding the thermodynamics of quadruplex unfolding can contribute to better design of quadruplex-interactive compounds, and improve the algorithms used to evaluate quadruplex formation and stability (36).

Water needs to be taken into account as a ligand for complete physical understanding of the macromolecule stability and conformation (37). The particular conformation of duplex, triplex, or quadruplex DNA depends directly on the degree of hydration, so changes in the water activity can shift the equilibrium to favor one conformation over others (36, 38, 39). It has also been shown that small osmolytes (e.g., ethylene glycol, glycerol, acetamide, sucrose) can decrease the water activity and destabilize duplex and triplex DNA (40). On the other hand, decreased water activity stabilizes quadruplexes (26, 41, 42), which indicates that quadruplexes are less hydrated in their folded form, and that water is taken up on their unfolding. Also, conformational changes of G-quadruplexes can be observed when the water activity is decreased, as shown for the human telomeric quadruplex in potassium solutions (43).

It has been shown that folding of *Tel22* can be described as an equilibrium of a three-state process, which suggests the formation of a stable intermediate triplex structure in the pathway between fully unfolded structure and the intramolecular G-quadruplex structure (3). The goal of the present study is to develop web application that could be used to perform thermodynamic characterization of G-quadruplex unfolding and test it by describing thermodynamic behavior of *Tel22* unfolding. Web application uses spectroscopic and calorimetric data at different salt and co-solute concentrations as an input to calculate thermodynamic parameters of unfolding mechanism of *Tel22*. Our results show that the unfolding mechanism of *Tel22* can be explained by a three-state equilibrium model (Fig. 1) and that K^+^ ions are released during the unfolding, whereas water molecules are taken up.

**FIGURE 1.**
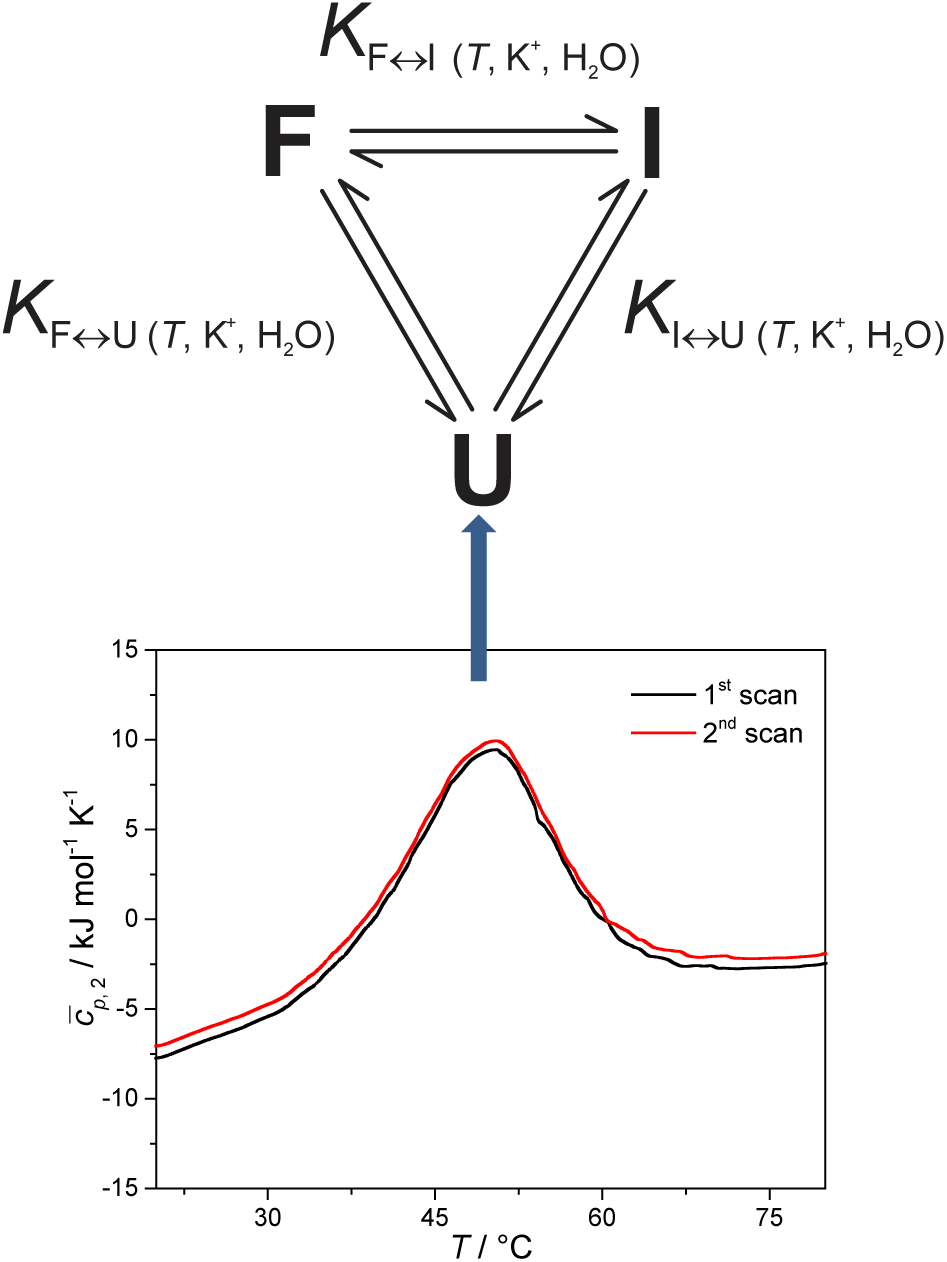
Proposed three-state equilibrium model for thermally induced folding/ unfolding of *Tel22* (top). Reproducible differential scanning calorimetry (DSC) heating thermograms confirm that the folding and unfolding of *Tel22* can be considered as an equilibrium process. The shape of the DSC thermograms indicate at least two thermally induced structural transitions (bottom).

## MATERIAL AND METHODS

### Sample preparation

*Tel22* is the oligonucleotide 5’-AGGGTTAGGGTTAGGGTTAGGG-3’, and it was obtained as HPLC pure from Midland Certified Reagent Company (USA). This DNA was first dissolved in water, and then extensively dialyzed (three buffer changes in 24 h) against 10 mM cacodylate buffer with 1 mM EDTA (pH 6.9) using dialysis tubing (Float-A-Lyser; Spectrum Laboratories, USA; M_w_ cut-off, 500-1000 Da). The DNA concentration in the buffer was determined spectrophotometrically at 25 °C. For the extinction coefficient of the single-stranded forms at 25 °C, a value of ε_260_ of 228.500 M^-1^cm^-1^ was used, which was obtained by extrapolation of the tabulated values (44) at 25 °C to 90 °C using procedures reported previously (45). The starting solution of the oligonucleotide was initially heated to 90 °C in an outer thermostat for 5 min, to make sure that all of the DNA was transformed into an unfolded form. This was then cooled to 5 °C (cooling rate, 0.5 °C/min), to allow the DNA to adopt quadruplex structure(s). This was then used in the study.

### Solution conditions

The buffers used consisted of 10 mM cacodylic acid (DSC, CD, UV) with 1 mM EDTA, with the addition of various concentrations of K^+^ ions and ethylene glycol. KOH was added to cacodylic acid to pH 6.9, and then KCl was added to obtain the desired of K^+^ ion concentrations (15, 55, 110 mM). To determine the effects of water activity on *Tel22* stability, ethylene glycol was added to the buffer with the lowest K^+^ ion concentration, to provide 3 M ethylene glycol.

### UV melting

The absorbance *versus* temperature profiles of the DNA samples were measured in a spectrophotometer (Cary 100 BIO UV/visible; Varian Inc.) that was equipped with a thermoelectric temperature controller, using cells of 1.0-cm path length. The melting of *Tel22* at a heating rate of 1.0 °C/min, followed by its formation at the cooling rate of 1.0 °C/min, were monitored at 260 nm and 293 nm, between 5 °C and 90 °C. Accurate concentrations were obtained at 25 °C from the melting curves monitored at 260 nm.

### Circular dichroism spectroscopy

The circular dichroism (CD) spectra of the G-quadruplexes were determined as a function of temperature in a CD spectropolarimeter (62A DS; AVIV) equipped with a thermoelectric temperature controller. The CD spectra of the samples (*c*_DNA_ ≈ 0.15 mM, in single strands) were collected between 215 nm and 320 nm in a 0.25-mm cuvette with a signal averaging time of 3 s and a bandwidth of 5 nm. The CD spectra were measured in the 5 °C to 90 °C temperature range, and at temperature intervals of 3 °C.

### Differential scanning calorimetry

Differential scanning calorimetry was performed using a Nano DSC instrument (TA Instruments, USA). The *Tel22* concentration used in the DSC studies was *c*_DNA_ ≈ 0.15 mM, in single strands. Cyclic DSC measurements were performed at the heating and cooling rates of 0.5, 1.0, and 2.0 °C/min. The measured temperature interval was between 1 °C and 90 °C. The corresponding baseline (buffer–buffer) scans were subtracted from the unfolding/ folding scans of the samples, and normalized to 1 mole of G-quadruplex in single strands, to obtain the partial molar heat capacity of the DNA, 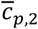, as a function of temperature. These data were analyzed in a model-dependent way by fitting the model function based on the reversible monomolecular three-state model (Fig. 1) to the corresponding experimental thermodynamic functions (see below).

### Thermodynamics of Tel22 folding/ unfolding

For each transition step i↔j (i↔j = F↔I, I↔U, F↔U) in the proposed folding/ unfolding mechanism (Fig. 1), the temperature-, cation-concentration-, and water-activity-dependent apparent equilibrium constant was calculated as given in Eq. 1:

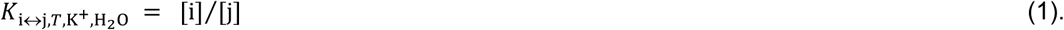

The real equilibrium constant is cation and water independent, and can be calculated as in Eq. 2:

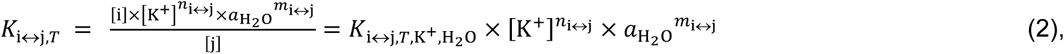

where *n*_i↔j_ is the number of K^+^ ions and *m*_i↔j_ is the number of water molecules that are released or taken up in the transition step i↔j, [K+] is the equilibrium concentration of unbound K^+^, normalized to a 1 M concentration in the reference state, and 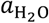 is the water activity. The water activity is given by Eq. 3:

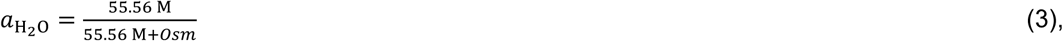

where *Osm* is the osmotic concentration of ethylene glycol (*[EG]*), which can be calculated according to Eq. 4, as proposed by Olsen et al. (26):

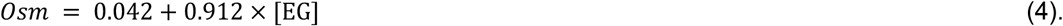

Each conformational transition of i↔j can be described in terms of the Gibbs energy by rearranging Eq. 2 to Eq. 5:

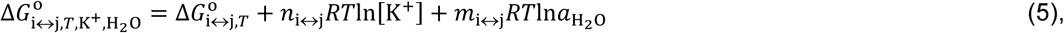

where 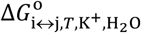 is the apparent, and 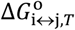 the true, thermodynamic Gibbs energy. Using the Gibbs-Helmholtz relationship, Kirchhoff’s law, and the reference temperature *T*_o_ of 298.15 K, the apparent Gibbs energy can be expressed as in Eq. 6:

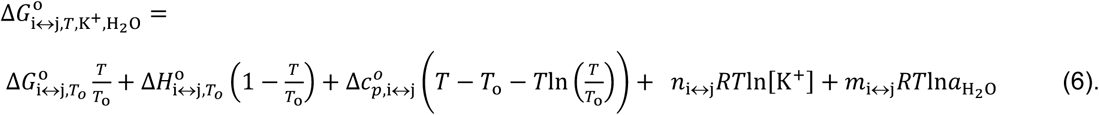

Each equilibrium constant that appears in the proposed model (see Fig. 1) can be expressed as in Eq. 7:

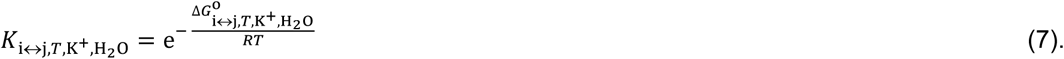

It can be seen from Eq. 2, 6, and 7 that to specify the population of any species in a single i↔j transition step, five parameters need to be calculated: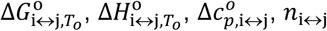, and *m*_i↔j_. As all of these parameters are state dependent, only 10 parameters (rather than 15) are needed to calculate the population of species F, I, and U in the solution at any *T*, K^+^, and ethylene glycol concentration. The calculated population of species i can be expressed as the molar ratio given in Eq. 8:

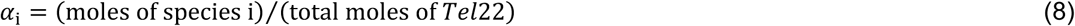

The calculated values of *α*_F_, *α*_I_, and *α*_U_ can be used to express the normalized CD signal, *f*, thus obtaining the CD model function shown in Eq. 9 (3, 46):

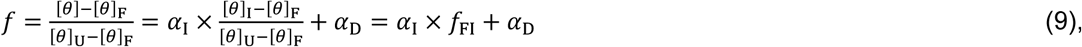

for which the [*θ*]_F_ and [*θ*]_U_ values are obtained as linear extrapolations of the baselines that describe folded and unfolded states over the measured temperature range. In Eq. 9, *f*_FI_ is considered as a temperature-independent normalization coefficient, which was treated in the model analysis as an adjustable parameter. At the same time, the values of *α*_F_, *α*_I_, and *α*_U_ can be used to express the excess heat capacity, which thus obtains the DSC model function of Eq. 10 (3, 46):

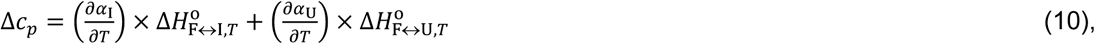

where 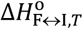 and 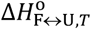 are the standard enthalpies for the interconversion of F to I and the unfolding of F to U, respectively. Δ*c*_*p*_ is obtained experimentally by subtracting the intrinsic heat capacity, 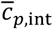, approximated by the second-order polynomial on *T* from the measured partial molar heat capacity of the DNA,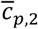. The global fitting of the DSC-based and CD-based model functions to the experimental DSC and CD data measured at different K^+^ concentrations and water activities was based on the nonlinear Levenberg-Marquardt *χ*^2^ regression procedure.

### Web application to calculate and fine-tune the model parameters

We developed a web application that enables the user to calculate and fine-tune the model parameters in an interactive and visual fashion. Web application is accessible through the following link: http://ltpo2.fri1.uni-lj.si/dna_old_formula/ Sample screenshots of the application are shown in Supplementary. A typical case-use scenario is as follows:

- The user uploads an arbitrary number of DSC datasets in the form of a CSV file. For each dataset, the file should specify the concentration of K^+^ ions, the concentration of ethylene glycol, and a series of temperatures – Δ*c*_p_ pairs (Fig. S1).
- Optionally, the user can upload an arbitrary number of spectroscopic datasets in a form similar to the DSC data (Fig. S2).
- The user now provides the initial values of the thermodynamic parameters and triggers the automated optimization procedure. The procedure computes the optimized values of the parameters using the Levenberg-Marquardt algorithm, then plots the calculated model function and the measurement function for each DSC dataset and spectroscopic dataset, and optionally calculates the error interval for each parameter using the bootstrapping technique. The user can save the optimized values of the parameters, revert to the old values if desired (either collectively or for each parameter individually), or fine-tune the parameters and continuously monitor the model and measurement function plots (Fig. S3 and Fig. S4).
- The user can control the optimization process by providing the optional upper and lower limits for the individual thermodynamic parameters, and by specifying which, if any, of the parameters should be kept intact during the optimization procedure. At the same time, the application makes it possible to specify the number of function evaluations in the Levenberg-Marquardt algorithm, and the number of iterations in the error estimation procedure (Supplementary Fig. S5).

The back-end of our web application was programmed in Python, using the Flask framework and the NumPy and SciPy libraries. The front-end was developed using a combination of various web technologies, including jQuery, Bootstrap, Highcharts, and MathJax.

## RESULTS AND DISCUSSION

Building an application for thermodynamic characterization of DNA unfolding is a monumental task, primarily because of the variety of mathematical models that can be used. As a first step, a proof of concept was created here to test the technical viability of an application, and to determine whether the target audience will like our idea. Thus, the web application only offers a non-customizable three-state equilibrium model to date. The model was selected because it can be used to describe thermally induced conformational transitions of *Tel22* (3). To obtain the data feed for our web application, several experiments were performed.

### Equilibrium model of *Tel22* unfolding

To check whether equilibrium models can be used to describe thermally induced conformational transitions of *Tel22*, CD spectra and DSC thermograms were recorded at different heating and cooling rates. These data were reproducible, which thus confirmed that the folding and unfolding processes of *Tel22* in the presence of K^+^ ions can be described as an equilibrium process. Furthermore, the melting profile of the *Tel22* thermal unfolding monitored by DSC, CD and UV showed biphasic behavior. The simplest model that can be used to describe the biphasic behavior of this *Tel22* thermal unfolding is illustrated in Fig. 1. The model incorporates three distinguishable structural forms and transitions of *Tel22* that are dependent on temperature and K^+^ and ethylene glycol concentrations. The model also allows the thermally induced unfolding of *Tel22* to follow two paths – the transition of structure F into structure I, followed by the transition into the unfolded structure U, and the direct transition from structure F to the unfolded structure U.

### Using CD spectroscopy to probe *Tel22 structure*

The CD spectra of *Tel22* in 15 mM and 110 mM K^+^ solutions and in 15 mM K^+^ solutions with 3 M ethylene glycol are shown in Fig. 2. CD is a very useful experimental technique for characterization of G-quadruplexes. The shape of the CD spectra allows us to predict the DNA strand orientation, and to distinguish between parallel-stranded G-quadruplexes (positive peak at 260 nm; negative peak at 240 nm) and antiparallel-stranded G-quadruplexes (positive peak at 295 nm; negative peak at 265 nm). The CD spectra of *Tel22* showed a negative band at ∼240 nm, a plateau at ∼250 nm, a shoulder at ∼270 nm, and a strong positive band at ∼290 nm. This shape of the CD spectra is characteristic of hybrid type G-quadruplexes (28, 47). Furthermore, the complex shapes of these CD spectra suggest that *Tel22* in K^+^ solution is a mixture of at least two hybrid-type G-quadruplex structures (36). As can be seen from Fig. 2, the position and height of these peaks depend on the K^+^ and ethylene glycol concentrations. Increased K^+^ ions concentration resulted in stronger CD spectra at an unchanged *Tel22* concentration, which suggests more densely packed G-quadruplexes due to stronger K^+^ electrostatic screening of the negatively charged phosphate groups. Decreased water activity resulted in a stronger negative peak at 240 nm, a drop in the signal at 250 nm, an unchanged shoulder at 270 nm, and a stronger positive peak at 290 nm. These changes are subtle, and they might reflect a shift in the population of the hybrid *Tel22* conformations upon addition of ethylene glycol (36, 48).

**FIGURE 2.**
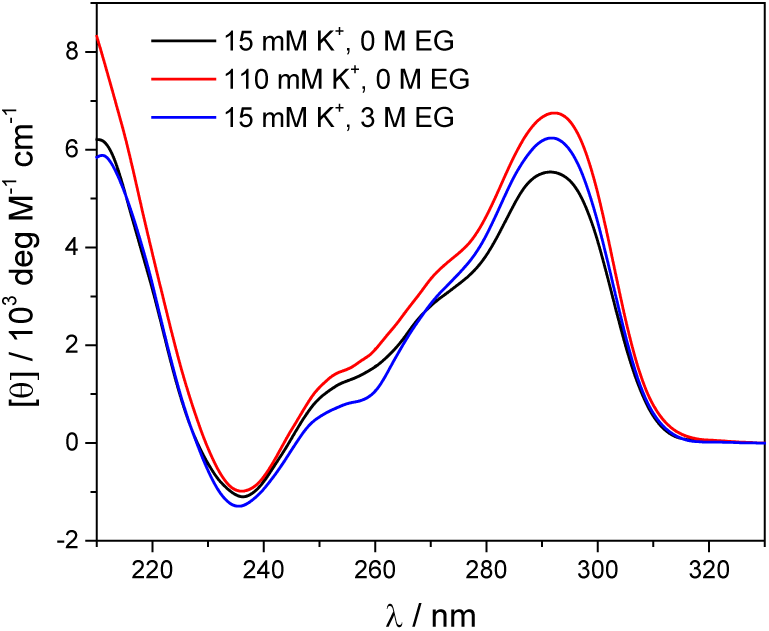
Circular dichroism spectra of *Tel22* at 5 °C in the presence of 15 mM and 110 mM K^+^ ions, and in the presence of 15 mM K^+^ ions and 3 M ethylene glycol, in cacodylate buffer, pH 6.9.

The melting profile of the *Tel22* quadruplex (*c* = 0.15 mM) was characterized in the presence of different K^+^ and ethylene glycol concentrations by monitoring the changes in ellipticity with increased temperature. Fig. 3 shows a representative recording of the CD spectra at different temperatures (c (K^+^) = 15 mM). The plot of ellipticity at 292 nm *versus* temperature yielded melting curves with melting temperatures, *T*_m_, that depended on the K^+^ and ethylene glycol concentrations. Increased K^+^ and ethylene glycol concentrations shifted *T*_m_ to higher values, thus demonstrating the stabilizing effects of K^+^ ions and ethylene glycol on these G-quadruplexes (Fig. 3). This effect was also confirmed by the UV-melting curves (data not shown), and it demonstrates the importance of the electrostatic interactions between the cationic metal ion and the anionic phosphate group for the stability of these G-quadruplexes. Additionally, quadruplex formation involves specific binding of certain metal ions, and their coordination with the guanine carbonyl oxygen atoms. For this coordination to take place, the metal ions must shed their water of hydration. As ethylene glycol not only decreases the water activity, but also the bulk dielectric constant, this allows the metal ions to shed their waters of hydration more easily, and thus increases the stability of the cation-quadruplex complex (48, 49).

**FIGURE 3.**
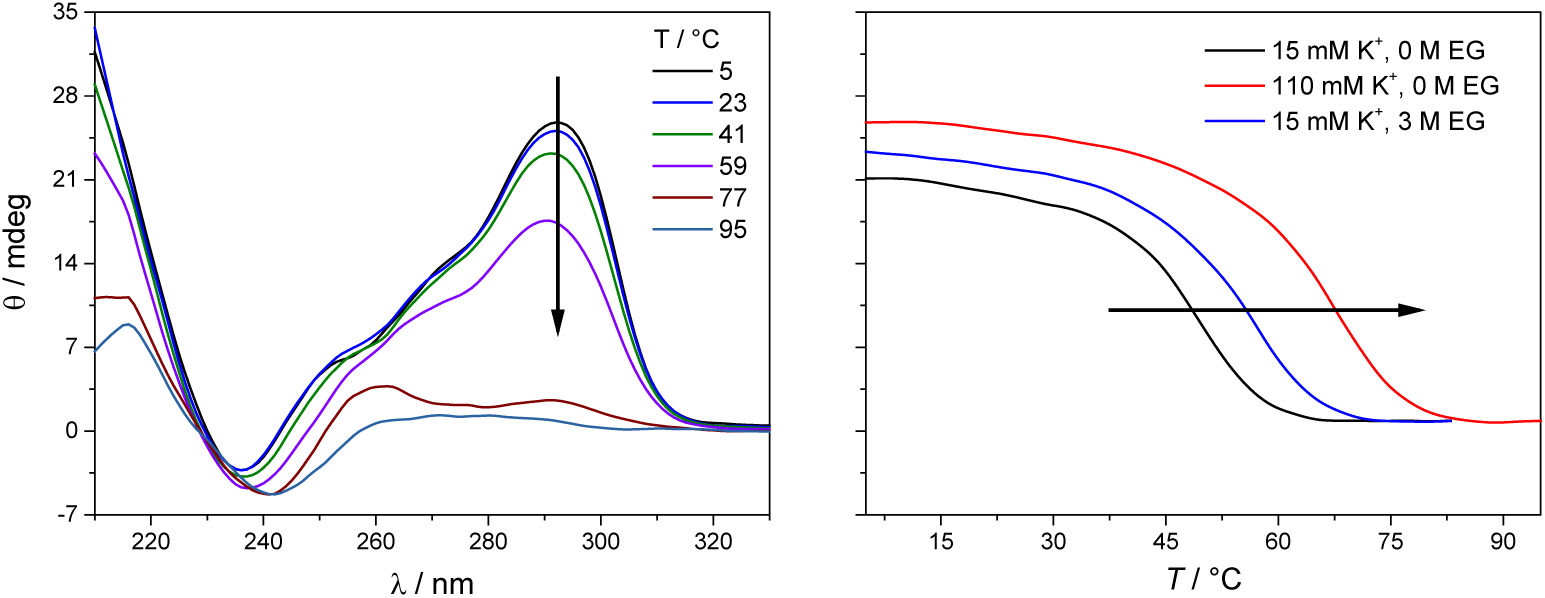
Temperature dependence of the circular dichroism (CD) spectra of *Tel22* quadruplexes in the presence of 15 mM K^+^ (right), and the CD melting curves obtained from the CD spectra at increasing temperatures at *λ* = 292 nm in the presence of 15 mM, 110 mM K^+^, and 3 M ethylene glycol (left). The concentration of *Tel22* was 0.15 mM, the path length of the cuvette was 0.25 mm, and the cacodylate buffer was at pH 6.9. The black arrows indicate the directionality of the changes in ellipticity with increasing temperatures (left) and the changes in ellipticity at 292 nm with increasing K^+^/ ethylene glycol concentrations.

### Global fitting of DSC and CD data

To obtain the complete thermodynamic profile of the thermally induced *Tel22* unfolding, the DSC melting curves were measured at different salt and ethylene glycol concentrations. On the basis of several specific examples of DNA and protein unfolding, it has been shown that the global thermodynamic analysis of experimental data obtained from several experimental techniques is essential for critical evaluation of model mechanisms (46). Thus, we uploaded experimental CD and DSC data to the web application to perform global fitting of the model functions (Eqs (9), (10)). NanoAnalyse was used to subtract the baselines before uploading the DSC curves to the web application. Although it is recommended to allow baseline correction for each species involved in the unfolding process during a fitting process (36), this option is not yet available for the web application, but it will be added in following versions. The CD melting curves were also normalized prior to being uploaded, but manual/automatic adjustments of the normalization procedures will be implemented in future versions of the web application. Next, the thermodynamic parameters obtained by Bončina et al. (3) were uploaded from a file. These parameters were used as initial values in fitting procedure and bootstrapping was used to calculate the errors.

Fig. 4 shows the comparison of model-based and measured CD melting and DSC heating curves and the thermodynamic parameters obtained by web application are shown in Table 1. When comparing these results to values obtained by Bončina et al. (3) some differences can be found, but the main conclusions for the overall unfolding process F↔U of *Tel22* remain the same: (i) 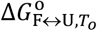 was positive and relatively small, due to the enthalpy–entropy compensation; (ii) the enthalpy change was ∼200 kJ mol^-1^, which is comparable to the literature data (6, 50); (iii) the heat capacity change was positive and substantial 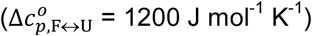; and (iv) K^+^ ions were released upon unfolding of *Tel22*.

**TABLE 1.** Thermodynamic profile of *Tel22* unfolding in the presence of K^+^ ions, obtained through global analysis of the differential scanning calorimetry and circular dichroism data based on the three-state model.

| Parameter | F ↔ I | I ↔ U | F ↔ U |
| --- | --- | --- | --- |
| $\Delta G_{i \leftrightarrow j, T_0}^0$ / (kJ mol<sup>-1</sup>) | 12.11 ± 0.54 | 27.52 ± 0.54 | 39.6 ± 1.1 |
| $\Delta H_{i \leftrightarrow j, T_0}^0$ / (kJ mol<sup>-1</sup>) | 89.4 ± 2.8 | 114.6 ± 3.4 | 204.0 ± 6.2 |
| $T \Delta S_{i \leftrightarrow j, T_0}^0$ / (kJ mol<sup>-1</sup>) | 77.3 ± 3.6 | 87.1 ± 3.9 | 164.4 ± 7.5 |
| $\Delta c_{p, i \leftrightarrow j}^0$ / (J mol<sup>-1</sup> K<sup>-1</sup>) | -315 ± 67 | 1510 ± 50 | 1200 ± 120 |
| $n_{i \leftrightarrow j}$ | 0.43 ± 0.04 | 1.88 ± 0.04 | 2.31 ± 0.08 |
| $m_{i \leftrightarrow j}$ | - 3.6 ± 3.0 | - 30.8 ± 2.6 | - 34.4 ± 5.6 |
| $f_{i \leftrightarrow j}$ | 0.40 ± 0.03 | / | / |

**Fig. 4.**
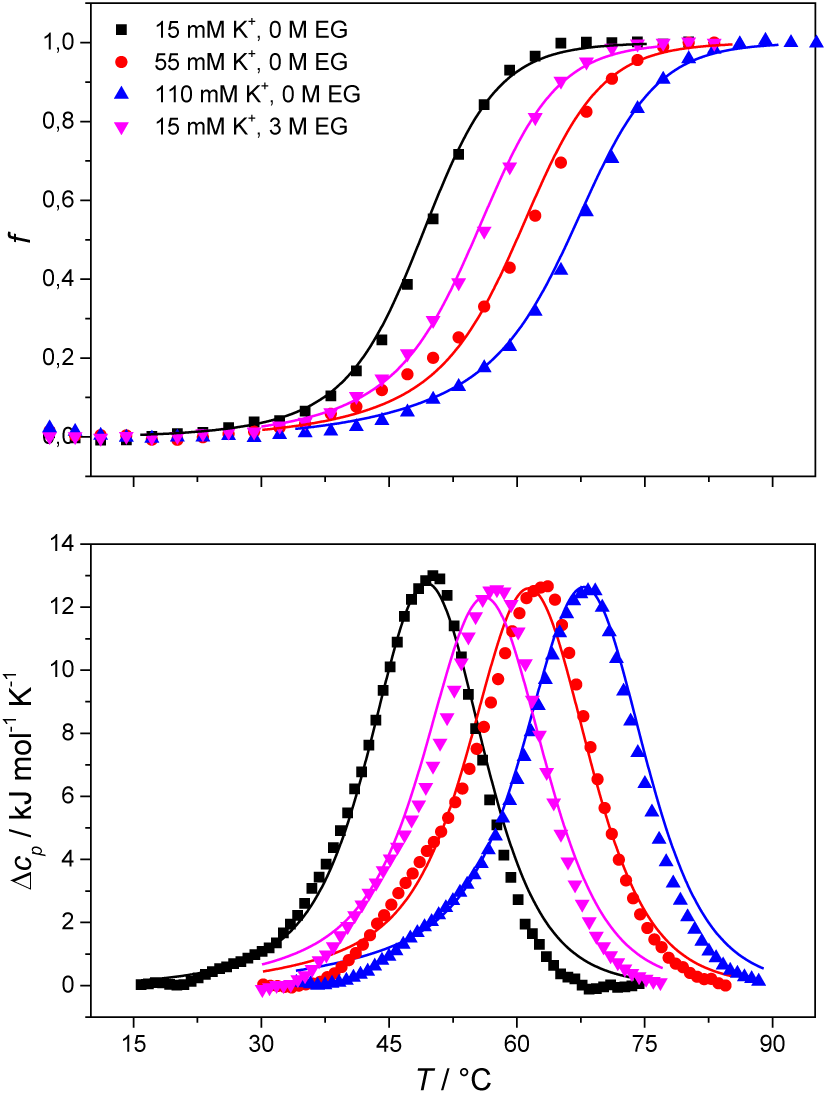
Comparison of the measured (points) and model-based (lines) normalized circular dichroism (CD) melting curves (top) and differential scanning calorimetry (DSC) heating curves (bottom) of *Tel22*, prepared in 15 mM, 55 mM, and 110 mM K^+^, and in 15 mM K^+^ and 3 M ethylene glycol, in cacodylic buffer, pH 6.9. The CD melting curves were recorded at 292 nm, the path length of the cuvette was 0.25 mm, and the concentration of *Tel22* used in the CD and DSC experiments was 0.15 mM.

### Water behavior during thermally induced unfolding of *Tel22*

The proposed model allowed us to study the effects of water activity on the thermally induced unfolding, and to calculate the number of water molecules that were released or taken up during the unfolding. Fig. 4 shows that addition of ethylene glycol (i.e., a decrease in the water activity) resulted in an increased *T*_m_, which is in favor of stabilizing effects of ethylene glycol on these *Tel22* quadruplexes. We obtained negative *m*_i↔j_ values for the unfolding of the G-quadruplexes (Table 1), which shows the uptake of water molecules. Specifically, per molecule of *Tel22*, we determined the water uptake of four molecules of H_2_O during the F ↔ I transition, and the water uptake of 31 molecules of H_2_O during the I ↔ U transition. According to Olsen et al. (26), these values can be attributed to several contributions to the unfolding of a quadruplex: (i) take up of structural water by the unfolded state; (ii) release of electrostricted water by the quadruplex, due to its lower charge density in the unfolded state; (iii) and take up of electrostricted water by the K^+^ ions upon their release from the G-quadruplex core. All three of these contributions will influence the heat capacity change – the uptake of structural water and the release of electrostricted water by the unfolded state will increase the heat capacity, while the uptake of electrostricted water by K^+^ will decrease the heat capacity. As the overall unfolding process F↔U is accompanied by a positive heat capacity change 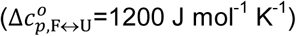 and a net uptake of water molecules (*m*_F↔U_=-35 molecules of H_2_O/molecule of *Tel22*), we can speculate that the main event responsible for the redistribution of water molecules during the F ↔ U transition is the exposure of a large number of thymines and other hydrophobic groups to the water. This is consistent with the calculated hydration heat capacity of nucleic-acid constituents, which shows that the melting of duplex DNA is accompanied by an increase in heat capacity (51). The number of water molecules that are taken up during the unfolding of the K^+^-stabilized *Tel22* (35 molecules of H_2_O/molecule of *Tel22*) is of the same order of magnitude as the number of water molecules that have been reported to be taken up during the unfolding of Na^+^-stabilized *Tel22* (100 molecules of H_2_O/molecule of *Tel22*) (52). The difference between the number of water molecules being taken up during this unfolding of K^+^- and Na^+^-stabilized *Tel22* (i.e., 35 *vs*. 100 water molecules) can be in part attributed to the different types of loops present in the hybrid (e.g., propeller-type, lateral, diagonal) and in the antiparallel (e.g., lateral, diagonal) structures. Namely, different types of loops can lead to greater exposure of nonpolar nucleotides to the surrounding water, which will in turn decrease the number of water molecules taken up during the unfolding of *Tel22*. The difference in the water behaviors between K^+^- and Na^+^-stabilized *Tel22* can also be attributed to the different free energies of K^+^ and Na^+^ hydration (53). As the Na^+^ cation has more negative hydration free energy, it is more effective at disrupting the water structure than K^+^. The number of water molecules taken up in the first and second transitions are significantly different, and correlate with the corresponding heat capacity changes. Table 1 shows that the uptake of the four molecules of water during the F ↔ I transition is associated with a lower heat capacity change 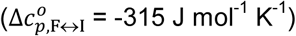 than the uptake of the 31 molecules of water during the I↔U transition, which is accompanied by a higher heat capacity change 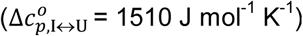. The differences between the first and second transitions are not limited to the water uptake and heat capacity change, but also extend to the other thermodynamic parameters that are characteristic for such individual transition steps (Table 1). Overall, we can conclude that the second transition is accompanied by greater changes in the thermodynamic parameters, which suggests that this involves more dramatic structural changes than the first transition. These results support the previously described *Tel22* unfolding mechanism, where intermediate state I was believed to be a triplex (3, 36). Partial unfolding of *Tel22* as such a triplex structure exposes only one G-quadruplex strand to the water, whereas the unfolding of the triplex structures itself exposes the remaining three G-quadruplex strands to water, which provides a large number of thymines and other hydrophobic groups. Consequently, the first transition (F ↔ I) is accompanied by a smaller heat capacity change than the second transition (I ↔ U).

### Temperature, potassium and ethylene glycol dependence of *Tel22* quadruplex species

The model analysis described here also provides the speciation diagrams at different K^+^ and ethylene glycol concentrations, as shown in Fig. 5. Comparisons of these curves that were calculated for all of the quadruplex species in the solutions with the corresponding measured DSC thermograms (Fig. 4) shows that the interconversion (F ↔ I) starts at relatively low temperatures (∼15 °C for 15 mM K^+^, ∼30 °C for 110 mM K^+^). It also shows that higher concentrations of potassium ions and addition of ethylene glycol not only shifts unfolding temperatures of *Tel22* to higher values but also increases intermediate concentration. The observed increase in concentration shows that intermediates are stabilized by higher K^+^ and EG concentration. This behavior suggests intermediates must exhibit several structural similarities with G-quadruplexes thus supporting our previous conclusion based on calculated thermodynamic parameters (Table 1) describing intermediate state as a G-triplex structure.

**FIGURE 5.**
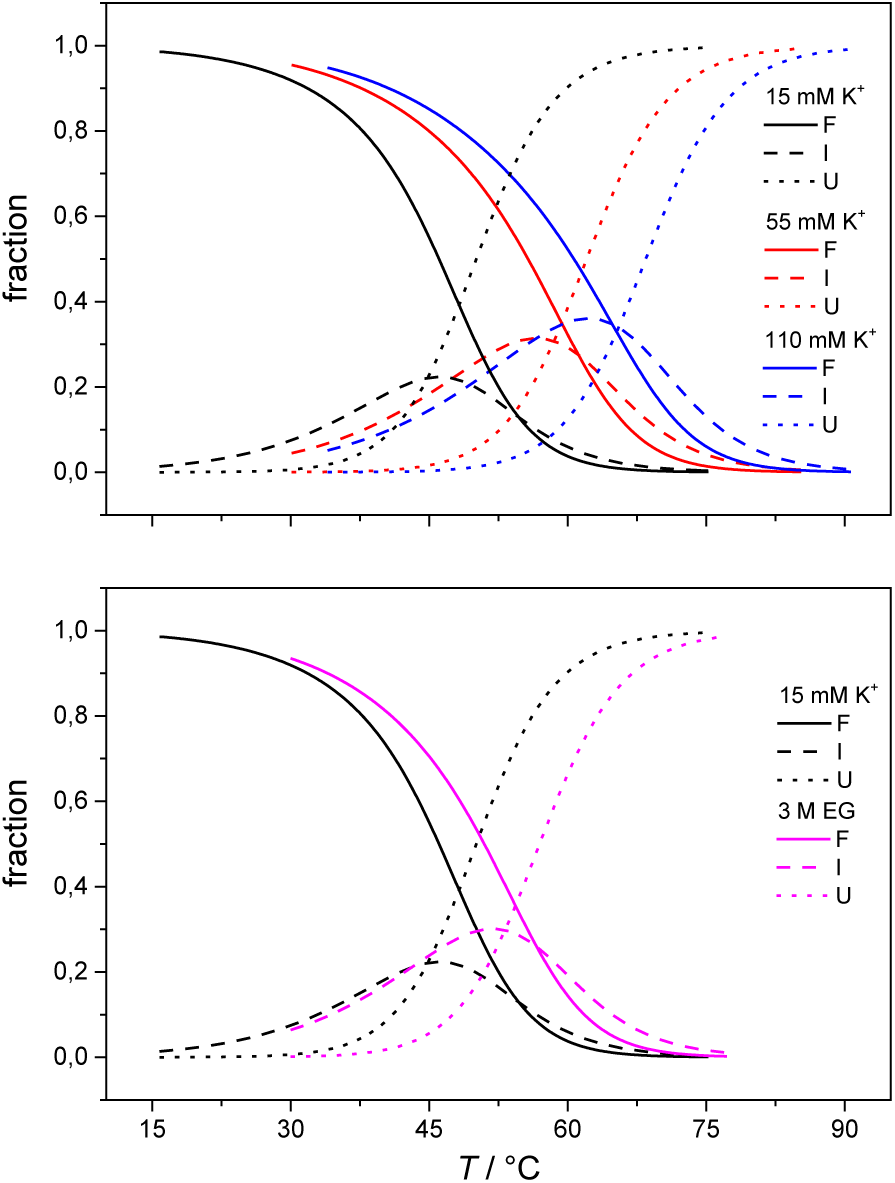
The model-based fractions of species F, I, and U as a function of *T* at the different concentrations of K^+^ ions (top) and ethylene glycol (bottom), determined using the best fit parameters reported in Table 1.

## CONCLUSION

The importance of thermodynamic parameters that determine the stabilities and interactions of biological macromolecules has contributed to an increase in the number of laboratories that are now equipped with sensitive microcalorimeters. Nowadays, thermodynamic data collection is comprehensive and routine, although thermodynamic data interpretation is still inadequate. This is partly because the general-purpose commercial software fails to provide tools to describe the behaviors of several unique biological systems, which leaves scientist with no other options but to develop their own software solutions.

As not everybody has the skills or time to build such tailor-made software, we are developing a web application that will be accessible to everybody, and will allow relatively easy thermodynamic analysis of calorimetric and spectroscopic data. This application can be used as a research and/or teaching tool and it will allow comparisons to be made of thermodynamic parameters obtained in different laboratories. As a proof of concept, we have tested the web application with experimental data obtained from monitoring thermal folding and unfolding of *Tel22* in the presence of different concentrations of K^+^ ions and ethylene glycol. We have thus described here the unfolding of *Tel22* according to a three-state equilibrium model (Fig. 1), and we have obtained the thermodynamic profile. The unfolding of *Tel22* was associated with positive changes in the free energy 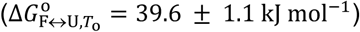, the enthalpy 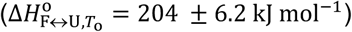, and the heat capacity 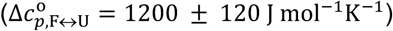, along with the release of K^+^ ions (*n*_F↔U_ = 2.3 ± 0.1) and the uptake of water molecules (*m*_F↔U_ = −34.4 ± 5.6).

Incorporating the effect of water activity in *Tel22* thermal unfolding model allowed our web application to calculate the number of uptaken water during transition of *Tel22* from folded to unfolded state. Comparing the number of uptaken water molecules to changes of heat capacity led us to conclusion that the main event responsible for redistribution of water molecules during *Tel22* unfolding is exposure of a large number of thymines and other hydrophobic groups to solvent. In addition, the comparison of thermodynamic parameters used to describe *Tel22* unfolding (Table 1) revealed big differences between first (F↔I) and second transitions (I↔U). Namely the second transition is accompanied by higher numbers suggesting that it involves more dramatic structural changes. This suggests that intermediate is structurally more similar to folded than unfolded state. Also speciation diagrams (Fig. 5) reveal that addition of ethylen glycol increases intermediate concentration during thermal unfolding. This behavior speaks in favor of interpreting intermediates as G-quadruplex related structures since stabilization of DNA sequences with decreased water activity is characteristic of G-quadruplexes. One possibility of such G-quadruplex related structure is G-triplex which was described in several publications as an important step in *Tel22* folding/unfolding mechanism.

The present study is important because it demonstrates the possibilities of performing thermodynamic analysis with specialized online tools. Our web application could be used to provide better understanding of driving forces responsible for the structural interconversion of G-quadruplex structures. These forces play a major role in the formation of G-quadruplexes and their physico-chemical properties and are undividedly related to the solution conditions (temperature, cation concentration/type and water activity). In the near future, we are planning to add several more models that can be used to describe folding and unfolding of biological macromolecules, the possibility to include experimental data from pressure perturbation calorimetry into this global fitting procedure and possibility to draw a phase diagram.

## AUTHOR CONTRIBUTIONS

I. P. wrote the manuscript, L. F: developed the web application, I. P. and S. S. designed and carried out whole research, N. P. U. contributed to discussion, data interpretation and writing of the manuscript.

## ACKNOWLEDGEMENT

We are grateful to Jurij Lah, Ph.D. for his helpful comments and suggestions and Chris Berrie, MA, MPhil, PhD for English language lecturing.

This work was supported in part by the Ministry of Higher Education, Science and Technology through the Slovenian Research Agency [grant numbers L7-8277, J4-8225, P4-0121].

